# Combined kinome inhibition states are predictive of cancer cell line sensitivity to kinase inhibitor combination therapies

**DOI:** 10.1101/2023.08.01.551346

**Authors:** Chinmaya U. Joisa, Kevin A. Chen, Samantha Beville, Timothy Stuhlmiller, Matthew E. Berginski, Denis Okumu, Brian T. Golitz, Gary L. Johnson, Shawn M. Gomez

## Abstract

Protein kinases are a primary focus in targeted therapy development for cancer, owing to their role as regulators in nearly all areas of cell life. Kinase inhibitors are one of the fastest growing drug classes in oncology, but resistance acquisition to kinase-targeting monotherapies is inevitable due to the dynamic and interconnected nature of the kinome in response to perturbation. Recent strategies targeting the kinome with combination therapies have shown promise, such as the approval of Trametinib and Dabrafenib in advanced melanoma, but similar empirical combination design for less characterized pathways remains a challenge. Computational combination screening is an attractive alternative, allowing in-silico screening prior to in-vitro or in-vivo testing of drastically fewer leads, increasing efficiency and effectiveness of drug development pipelines. In this work, we generate combined kinome inhibition states of 40,000 kinase inhibitor combinations from kinobeads-based kinome profiling across 64 doses. We then integrated these with baseline transcriptomics from CCLE to build robust machine learning models to predict cell line sensitivity from NCI-ALMANAC across nine cancer types, with model accuracy R^2^ ∼ 0.75-0.9 after feature selection using elastic-net regression. We further validated the model’s ability to extend to real-world examples by using the best-performing breast cancer model to generate predictions for kinase inhibitor combination sensitivity and synergy in a PDX-derived TNBC cell line and saw reasonable global accuracy in our experimental validation (R^2^ ∼ 0.7) as well as high accuracy in predicting synergy using four popular metrics (R^2^ ∼ 0.9). Additionally, the model was able to predict a highly synergistic combination of Trametinib (MEK inhibitor) and Omipalisib (PI3K inhibitor) for TNBC treatment, which incidentally was recently in phase I clinical trials for TNBC. Our choice of tree-based models over networks for greater interpretability also allowed us to further interrogate which specific kinases were highly predictive of cell sensitivity in each cancer type, and we saw confirmatory strong predictive power in the inhibition of MAPK, CDK, and STK kinases. Overall, these results suggest that kinome inhibition states of kinase inhibitor combinations are strongly predictive of cell line responses and have great potential for integration into computational drug screening pipelines. This approach may facilitate the identification of effective kinase inhibitor combinations and accelerate the development of novel cancer therapies, ultimately improving patient outcomes.

## 1. Introduction

Protein kinases, which serve as the primary conduits for information transfer within cells, are often implicated as key drivers in cancer development and have thus become a cornerstone in current targeted therapies [1]. The rapid expansion of kinase inhibitor therapies as an oncology drug class is exemplified by the FDA’s approval of nearly 60 such therapies over the past 20 years [2]. Despite their initial promise, kinase-targeting monotherapies frequently give rise to resistance [3], in part due to the dynamic nature of the kinase network, i.e., the “kinome,” which has been shown to reprogram and respond to the inhibition of single kinases by upregulating expression of partner pathways [4–6]. This necessitates the development of novel strategies to effectively target the kinome and harness the vast array of potential drug targets it offers.

One emerging strategy to counteract the dynamic acquisition of kinase inhibitor resistance involves the design of combination therapies, which perturb multiple targets with two or more drugs. These targets may be either known compensatory pathway partners, referred to as “horizontal pathway inhibition,” or multiple targets within the same pathway, known as “vertical pathway inhibition” [7]. This approach has recently gained traction with the FDA approval of the combination of trametinib and dabrafenib for treating advanced melanoma [8]. This combination therapy “vertically” targets both BRAF and MEK within the RAF-MEK-ERK (MAPK) pathway, demonstrating the potential effectiveness of this strategy. However, this method of empirical design of combination therapies is not feasible for less characterized kinase pathways, and the sheer number of possible combinations of potential kinase targets (2∼500) prevents comprehensive screening or drug design.

To circumvent this issue, computational screening offers an appealing alternative, enabling the prediction of effective drug combinations in-silico prior to testing a reduced pool of potential combinations in-vitro. This method not only potentially streamlines the drug development process but, when combined with patient-specific genomic profiling, can also enable personalized drug combination selection to potentially achieve resistance-proof responses in patients.

In recent years, a variety of computational approaches have been developed to predict combination therapy responses for cancer drug screening [9,10]. The majority of these methods primarily rely on drug structure characteristics and cancer-specific baseline genomic profiling to predict effective drug combinations, spurred by advancements in the high-throughput acquisition of these data types. For example, a high-dimensional tensor-based modeling strategy used similar data and achieved impressive accuracy (Overall R^2^ ∼ 0.8) in predicting response to combination therapies, validated experimentally [11]. This approach and others employ intricate neural network architectures that, while capable of producing high performing models, can be challenging to interpret, posing a barrier to the broader adoption and understanding of their underlying mechanisms. Tree-based machine learning models on the other hand, although simpler and sometimes less powerful, are generally considered interpretable depending on the type of data fed to them [12]. Notably, drug-protein interactions, which are intuitively central to the process of phenotype reversal, have been relatively underexplored in these computational approaches. In part, the minimal amount of drug-target information leveraged in current response prediction efforts is because of the sheer amount of data generated by genomics and molecular fingerprinting, generating thousands of features for each measurement, while drug target data has been generally sparse with only a few annotated targets per drug. However, recent advances in technology to profile the interactions of clinical drugs with all the members of the kinome represent an unprecedented ability to measure drug-target information across ∼500 proteins simultaneously in a quantitative manner [13,14]. The breadth, density, and ease of acquisition of this data, often measured at multiple dose points, is ideal for integration into machine learning models that can leverage diverse data types for drug response prediction.

Specifically, recent advances in proteomics techniques have facilitated the large-scale characterization of drug-kinase interactions, providing valuable information on the extent to which the entire kinome is inhibited by specific drugs or drug combinations. A landmark paper in 2017 used a mass spectrometry-based assay that used promiscuous kinase-binding compounds immobilized on beads to measure the binding competition between any given inhibitor and any given kinase (henceforth called the “kinobeads” assay) [15]. Using this assay, the kinome-wide binding profiles for ∼230 clinical kinase inhibitors at eight doses each were elucidated using cancer cell lysates, forming the largest in-cell drug-target binding database publicly available at this time. The data generated from these assays allow interrogation of how clinical and investigational drugs interact with the entire kinome on an unprecedented scale. By analyzing the degree of inhibition of all kinases simultaneously for a given inhibitor, we can treat this as characterizing the degree of departure from the “baseline kinome state”, thus moving through drug-induced alteration of multiple kinase activities to a new “kinome inhibition state”. Given the degree to which modulation of the kinome alters cellular state and downstream behavior, these baseline kinome states and kinome inhibition states can be directly connected to various measured cellular phenotypes. We have recently demonstrated this idea by showing that kinome inhibition state is significantly predictive of cancer cell responses to kinase inhibitor monotherapies when integrated with cancer-specific information, such as baseline transcriptomics, using tree-based machine learning models [16].

In this work, we show that by combining the inhibition states of two kinase inhibitors, we can generate a hypothetical “combined” inhibition state for an untested inhibitor combination. In this manner, we can rationally use all combinatorial kinome inhibition states to sample all possible kinase target combinations, hypothetically including all pathway partners. By integrating these inhibition states with cancer-specific baseline transcriptomics, we demonstrate that the combined inhibition state can predict the sensitivity of cancer cell lines to inhibitor combination treatments from the NCI-ALMANAC dataset using interpretable machine learning models. We further validate these models experimentally by examining novel inhibitor combinations in a PDX-derived triple-negative breast cancer (TNBC) cell line. By focusing on dual-inhibitor drug-kinase interactions combined with cancer-specific baseline genomic profiling, we can enhance computation combination drug screening pipelines with combinatorial kinase targeting. Furthermore, this approach lays the foundation for the rational design and a priori prediction of combination kinase inhibitor treatments for patients with the potential to ultimately reduce single kinase inhibitor resistance acquisition by prior rational targeting of partner pathways and associated kinases.

## 2. Results

### 2.1. Creating a Set of Combined Kinome Inhibition States Representing Current and Potential Kinase Inhibitor Combination Therapies

In this work, we have focused on a specific set of 200 kinase inhibitors characterized using the kinobeads assay [15]. These inhibitors were profiled in-cell for their interactions with ∼500 kinases and kinase-interacting proteins, across eight doses. From this data, as described previously (insert citation), we extracted monotherapy “kinome inhibition states”, denoting the degree to which they inhibit each kinase in the kinome at eight doses on a scale of 0-1 (0 is complete inhibition and 1 is no inhibition of a given kinase).

We next tested different methods to approximate the kinome inhibition state of a kinase inhibitor combination. Intuitively, this can be thought of as simply superimposing two individual monotherapy inhibition states, but for the few cases where different inhibitors target the same kinase, we have to find ways to accurately reflect the resulting effect on the kinome. Here, we tested combining monotherapy kinome inhibition state vectors through addition, multiplication, truncated multiplication (excluding kinase inhibition values >1). All three methods were compared for downstream model performance.

After combining the individual inhibition states, we were left with a dataset describing all possible pairwise combinations of ∼220 kinase inhibitors. These ∼45,000 combinations represent the kinome inhibition states of existing clinical therapies (example), therapies currently in clinical trials (example), as well as potential therapies. Together, they interrogate a search space that includes nearly every known kinase on the phylogenetic tree (Fig S1).

### 2.2. Connecting Inhibited Kinome States with Cancer Cell Line Combination Sensitivities

Next, we linked the data set describing kinase inhibitor combinations to their cell sensitivity phenotypes in the large-scale ALMANAC drug combination screen. The ALMANAC screen contains cell sensitivity data for 53 kinase inhibitor combinations, over ∼200 unique dose combinations for 45 cell lines across 9 cancer types. Additionally, previous high-throughput combination screens conducted in our lab in breast cancer offered data for 56 inhibitor combinations in four cell lines. Ideally, we would like exact matches between the dose at which kinome inhibition state is profiled and the dose at which cell sensitivity was measured. However, there are very few exact matches between the datasets. To overcome this, we found the nearest dose (at maximum differing by 1uM) at which kinome inhibition was profiled for each cell sensitivity measurement and connected the two datasets using these dose matches.

Additionally, we added cell line specific information to the dataset to complement the drug-specific kinome inhibition states. The CCLE database contains baseline transcriptomics data for ∼1500 cancer cell lines, and almost all of the cell lines included in our data set were represented. Using this, we further added baseline gene expression into the dataset, now containing kinase inhibitor combinations, their inhibition state of the kinome, the cell line sensitivity to their treatment, as well as that cell line’s baseline gene expression. In this way, the dataset connects the kinome inhibition states of inhibitor combinations to their cell sensitivity phenotypes.

The collected dataset represents a total of eight major cancer types, with the majority having ∼7 cell lines represented each, while breast cancer had the most representation (11 cell lines). To ensure that the machine learning model downstream could find cancer-specific linkages between the kinome and cell sensitivity, we split the dataset into eight individual cancer type datasets and conducted all modeling on each data split in parallel.

### 2.3. Elastic-Net Feature Selection Reveals Kinome Inhibition States to be Most Informative

In our collected dataset, kinome inhibition states and baseline gene expression together represent ∼20,000 variables or “features” that could affect the phenotype of cell sensitivity to kinase inhibitors. It is both practically prohibitive and ineffective to build models using all available features, and so keeping in mind computational efficiency we sought to filter down the dataset to include only the most informative features. To accomplish this “feature selection”, we built our machine learning pipeline starting with an elastic-net regression [17] model built against the outcome of cell sensitivity. This generated coefficients for each feature, with the absolute value of the feature coefficient directly proportional to its predictive value for the outcome. We ensured non-informative features were not included in modeling by only considering features with non-zero coefficients. We fit the model on the entire dataset to visualize a snapshot of the feature coefficients globally. This revealed overwhelmingly larger coefficients for kinome inhibition states compared to baseline gene expression (Fig 2a), thus indicating that kinome inhibition states were globally more informative for cell sensitivity prediction compared to baseline gene expression.

**Figure 1.**
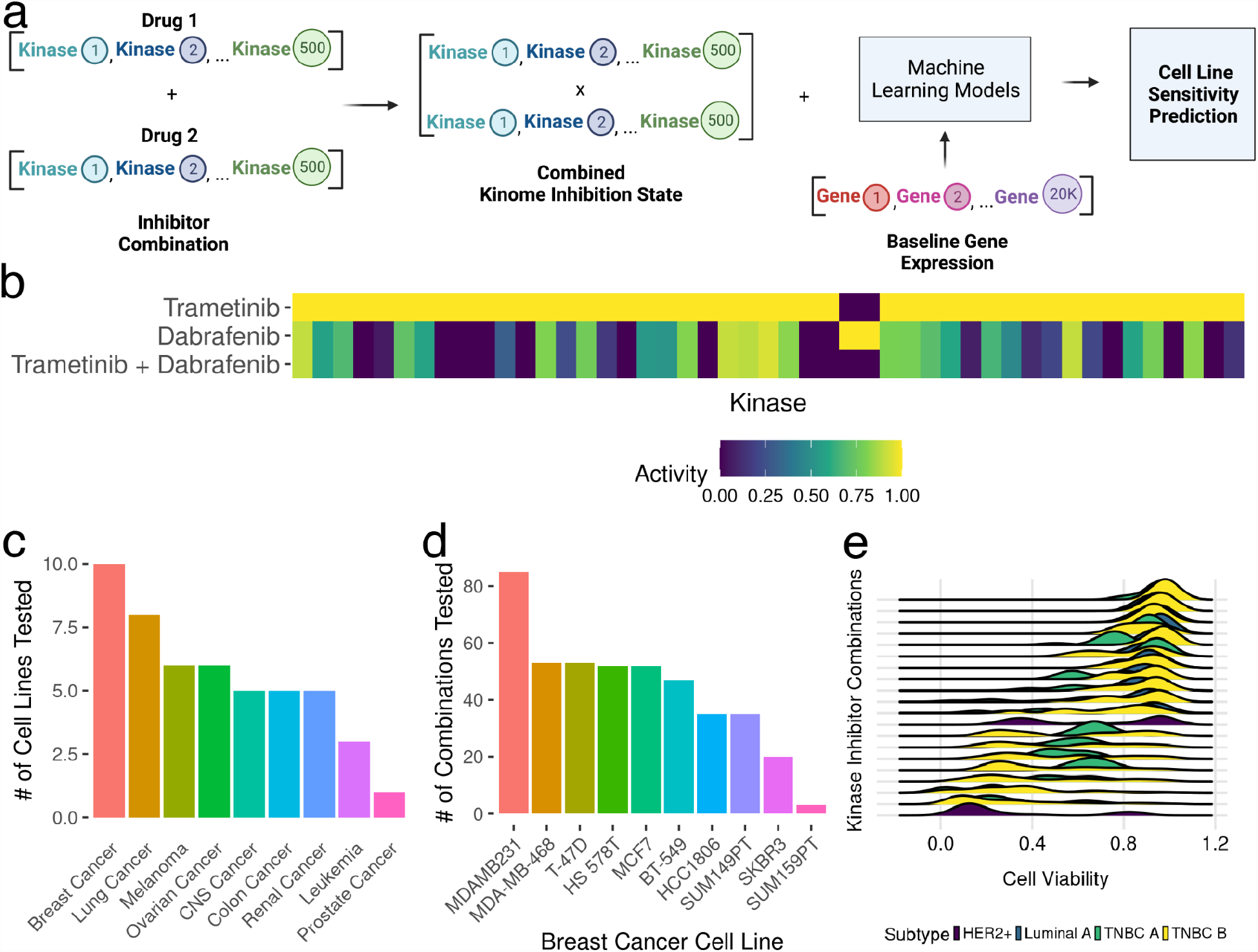
Kinome inhibition State Combination Modeling and Data Overview. (a) Schematic of modeling pipeline. (b) Heatmap showing the inhibition state of individual kinase inhibitors (row 1 and 2), and the hypothetical “combined” inhibition state for the two inhibitors (row 3) (c) Bar plot showing number of cell lines tested per cancer type in training data set (d) Bar plot showing number of unique combinations tested per cell line for the breast cancer subset of the training data set (e) Ridge plots showing cell viability (x-axis) variation for a random subset of different kinase inhibitor combinations (y-axis) in the NCI-ALMANAC data for breast cancer cell lines. Different breast cancer subtypes are represented with differing colors.

**Figure 2.**
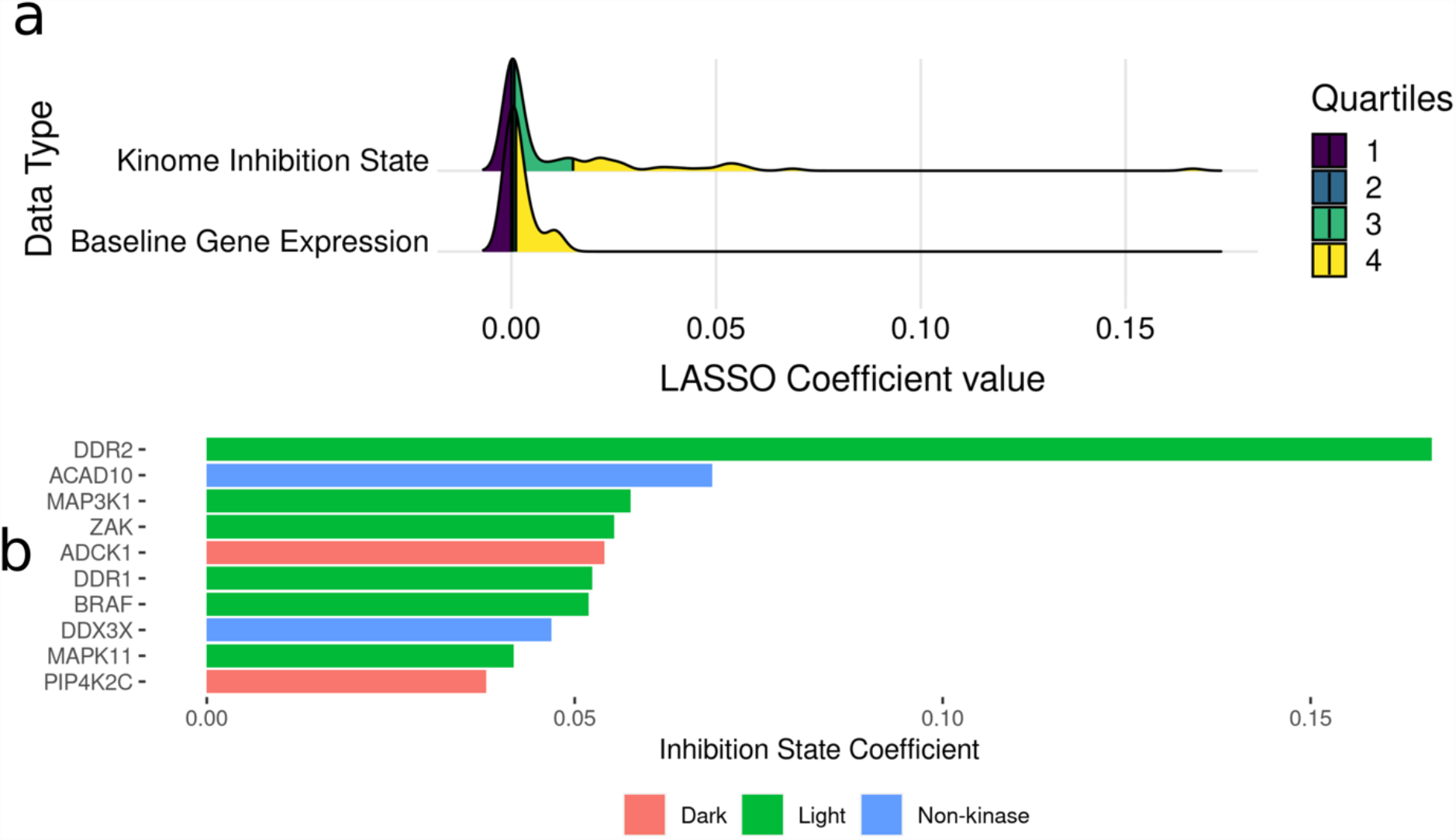
Feature Selection using an Elastic-net Regression Model against Cancer Cell Line Sensitivity. (a) Ridge plot showing the distribution of LASSO coefficient sizes as a metric for feature importance, for each feature type (b) Horizontal bar plot showing kinases with the largest elastic-net coefficient values, coloured by whether they are defined as “understudied” (Dark) or “well-characterized” (Light).

For downstream model building, the data set was split into a training and testing set five times (five-fold cross validation). For the training set data to not have any influence on the test set (to prevent data leakage), the elastic net model is fit on only the training data, and features are selected within each fold. Parameters for the elastic net model and hyperparameters for the tested model types were also tuned this way.

### 2.4. Machine Learning Models Can Predict Cancer Cell Line Sensitivity to Combination Therapies by Integrating Kinome Inhibition States and Baseline Transcriptomics

After data set preparation and feature selection, we built machine learning models that can predict cell sensitivity to kinase inhibitor combinations. For each cancer type, three machine learning model types were tested: random forest, boosted trees (xgboost) and deep neural networks. Xgboost performed the best for all cancer types, with type-specific performance largely dependent on abundance of data in the training set (Fig 3b). The most abundant cancer type (breast) had the best performing model with an R^2^ score of 0.93 (Fig 3b) while the lowest performing model was prostate cancer with R^2^ = 0.73. Given our previous lab experience with breast cancer, we chose the breast cancer model for downstream experiments and validation.

**Figure 3.**
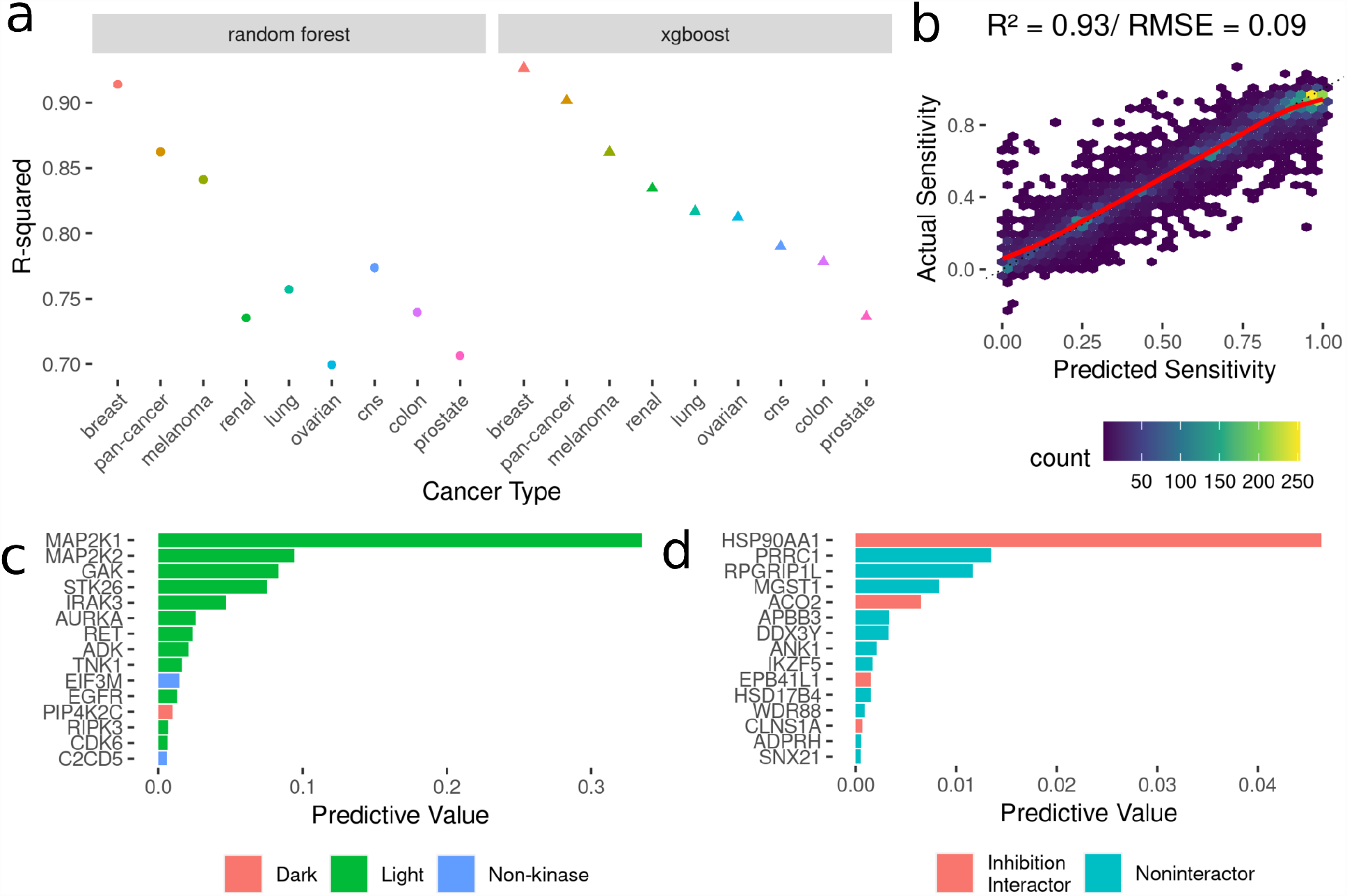
Development of Models to Predict Cancer Cell Line Sensitivities to Kinase Inhibitor Combination Therapies from Kinome Inhibition States. (a) Model performance metrics (R-squared) for Random Forest (dots) and XGBoost (triangles). (b) Scatter Plot of predicted sensitivity values from the best-performing model vs actual sensitivity values. The red line indicates a smooth fit through the data points. (c) Horizontal bar plot showing model importance of individual kinase inhibition states by importance values. (d) Horizontal bar plot showing model importance of individual baseline gene expression by importance values.

Additionally, since the best-performing model was tree-based gradient boosting, we were able to further analyze the model using computed feature importance to find the most informative features in the data set based on the feature importance metric. Similar to the feature selection output, we saw much higher feature importance scores overall for kinome inhibition states when compared to baseline gene expression, and several kinases implicated in breast cancer dysfunction had high importance scores, such as MAP2K1/2 and EGFR(Fig. 3c).

### 2.5. Experimental Validation of Model Predictions in a PDX-Derived Triple Negative Breast Cancer Cell Line was Successful

We demonstrated that machine learning models using the kinome inhibition states of combination therapies along with cell-specific baseline gene expression could robustly predict cell sensitivity in multiple cancer types. However, to see if these predictive models could extend to real-world experiments, we experimentally validated 35 kinase inhibitor combinations in a PDX-tumor derived cell line(Fig 4A).

**Figure 4.**
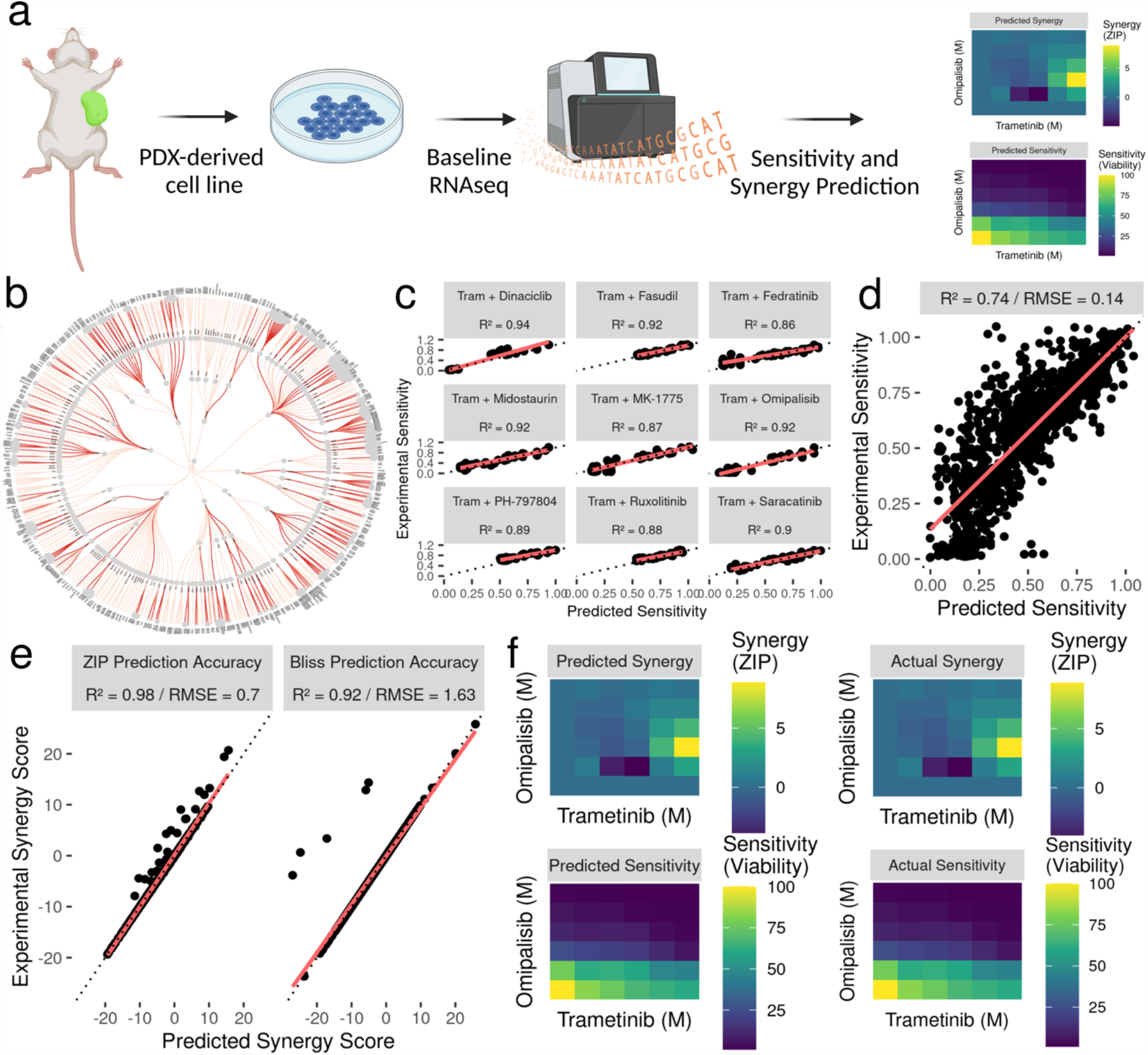
Experimental Validation of Model through a Trametinib Combination Screen in the WHIM12 Patient-Derived TNBC Cell Line. (a) Schematic showing experimental validation pipeline for the WHIM12 PDX-derived cell line. (b) Kinome phylogenetic map showing diversity of kinome space targeted (red = inhibited by a validated kinase inhibitor combination). (c) Grid of scatter plots showing accuracy of predicted vs experimental sensitivity to the top 9 tested combinations. For all scatter plots, the dashed line indicates where perfect predictions would lie and the red line shows a linear fit through the data. Quantitative accuracy is represented by the R-squared score. (d) Scatter plot showing the global accuracy of model predicted sensitivity compared to experimental sensitivity. (e) Grid of scatter plots showing accuracy of model predicted synergy scores compared to experimentally measured synergy scores across two metric types (ZIP, Bliss). (f) Grid of heatmap plots showing comparison of predicted vs experimentally measured sensitivity and synergy for the highly synergistic trametinib / omipalisib combination.

High-throughput cell line drug screens have been widely documented to suffer from a lack of reproducibility and poor translation to more complex samples like patient tumours. We sought to test whether our model of cell sensitivity in breast cancer, trained on 11 well-characterized immortalized cell lines, could effectively predict cell sensitivity in a PDX (Patient-Derived Xenograft) derived cell line. We chose the WHIM12 PDX-derived cell line, which was generated from a highly chemo-resistant TNBC tumor [18]. Previous experiments in the lab had conducted a drug combination screen in the WHIM12 cell line, out of which 35 kinase inhibitors were tested in combination with trametinib. Complementary baseline gene expression data was also generated through RNAseq. Using these in-house data, we were able to input the unseen WHIM12 gene expression into the trained model and predict the cell sensitivity outcomes of the conducted drug combination screen. We achieved robust prediction accuracy (Global R^2^ = 0.74 / RMSE = 0.14) in predicting exact cell viability in response to treatment with 35 kinase inhibitor combinations, across 64 dose combinations (Fig 4c, d).

### 2.6. Model Predictions Reveal Known Synergy in trametinib/omipalisib Combination

The model predictions in the WHIM12 cell line were further interrogated for potential synergy. We generated synergy scores for all 35 combinations at each of the 64 dose points using the R package SynergyFinder [19] based on four different metrics: Zero-Interaction Potency [10] (ZIP), Bliss Independence [20], Highest Single-Agent (HSA), and Loewe Additivity [21]. Additionally, we generated similar synergy scores using the actual experimental data generated for validation as a comparison. We found a high degree of similarity (Global R^2^ ∼ 0.94/ RMSE ∼ 0.5) between predicted and actual synergy, with trametinib + omipalisib as our most synergistic predicted combination, with a ZIP score of ∼8 at certain dose combinations (Fig 4e, f). This is significant as the model predictions were in a TNBC PDX-derived line, and the trametinib/omipalisib combination represents the popular strategy of simultaneously targeting the MAPK and PI3K pathways [22].

## 3. Discussion

Kinase inhibitors are one of the fastest growing drug classes for cancer therapy, with ∼62 FDA approved in total against neoplasms [2]. With 500 potential druggable targets, there is significant interest in decoding the spectrum of kinases targeted by current inhibitors, and streamlining the kinase inhibitor screening process. We have previously introduced [16,23,24] the idea that the full spectrum of a given inhibitor’s effect on the kinome as measured by recent advances in kinobead-competition/MS technology [15] can be represented as a “kinome inhibition state”, i.e. a vector representing the effect of a given inhibitor on the kinome as a whole.

In this work, we have extended this idea to represent the kinome inhibition state of a combination of inhibitors, using a simple multiplicative probability model to “combine” the inhibition states of two given kinase inhibitors. By generating these “combined” inhibition states, we can vastly expand the search space targeted by inhibitor monotherapies, sampling all possible combinations of currently available therapies. To accomplish this, we used publicly available drug-kinome interaction data to generate snapshots of the combined effect of a combination therapy on the protein kinome. We then linked these kinome inhibition states of inhibitor combinations to cancer cell sensitivity phenotypes to combination treatment, creating a framework for predicting the efficacy of combination therapies in different cancer types.

We fit tree-based machine learning models as well as neural networks on this linked data set to robustly predict precise cancer cell line sensitivity and synergy for untested kinase inhibitor combinations therapies and validate those predictions in complex patient derived samples. gradient-boosted tree models were highly accurate across cancer types (R^2^ 0.75-0.93), comparable to two recent neural-network driven attempts to predict cell line response to drug combinations [9,11]. We chose to move forward with the highest performing breast cancer model for further validation. We chose to validate our model predictions in the PDX-dervied WHIM12 line, reasoning that PDX-derived cell lines retain many of the molecular and genetic features of the xenografted original tumors and offer a closer representation of the disease state compared to traditional cell lines. We were able to show that the models performed robustly on novel baseline gene expression data (Global Accurcacy R^2^ ∼0.74), representing its ability to extend to complex and clinical-adjacent samples compared to well-characterized cell line data.

One of the strengths of tree-based models compared to deep neural networks is that they are generally considered to be interpretable through feature importance computation [12,25]. Using this, we were able to investigate the “black box” and query which specific kinase inhibition states and baseline genes were most predictive of cell sensitivity. We found that for the best performing breast cancer model, the inhibition of the kinases MAP2K1/2 were the most informative by far. This is intuitive considering the most abundant kinase inhibitor in the dataset is the allosteric MEK inhibitor trametinib, but it must be noted that MEK inhibition is always only just one half of the kinome targeting in the combination. There has been increasing clinical interest recently in targeting the PI3K and MAPK pathways [22], and our lab has shown before that MEK1/2 inhibition in TNBC by trametinib induces widespread transcriptional adaptation, and that there is potential for clinical efficacy for complementary kinome targeting with trametinib [26]. Since our model’s sensitivity predictions can effectively simultaneously predict synergy, our top synergy prediction for breast cancer according to the ZIP metric was trametinib and omipalisib, which we were able to validate experimentally in the WHIM12 line. This indicates that from the training breast cancer screening data, the model was able to learn that targeting the complementary PI3K and MAPK pathways is effective and synergistic in TNBC cell lines.

Interestingly, the predicted high-synergy combination of trametinib/omipalisib was recently in phase I clinical trials for advanced solid tumors but failed due to patient intolerability [27]. This highlights some limitations of our modeling approach. Ideally, kinome inhibition state would be one of many different drug modalities included for response prediction, and we plan to further expand these models in the future by considering toxicity, drug structure and cancer-describing multi-omic data types not limited to baseline gene expression. Additionally, in this proof-of-concept study we utilized a simple multiplicative probability model to generate the “combined” inhibition state of two inhibitors on the kinome, by assuming that the inhibition of a given kinase is mutually exclusive from that of other kinases. We know that kinases function physiologically as part of complex signaling networks, and their inhibition may have downstream effects on other kinases and signaling pathways. To address this limitation, future models will incorporate more biologically representative schemes to hypothesize combined kinome inhibition states.

In summary, through this work we demonstrate the development of a framework for predicting the efficacy of combination therapies in different cancer types using just kinome-drug interactions and baseline gene expression. We use a multiplicative probability model to generate the “kinome inhibition state” of a combination of kinase inhibitors and link these states to cancer cell sensitivity phenotypes. First, we were able to show that a given combination therapy’s cancer-agnostic interaction with the kinome was far more informative than baseline genomics in predicting downstream response. This is intuitive fundamentally, as drug-protein interactions are the primary means of drug effect on physiology, but this type of data is still underutilized in computational screening approaches. We then used machine learning models to predict cell line sensitivity and synergy for untested kinase inhibitor combination therapies and validate those predictions in complex patient derived samples. We found that the inhibition of the kinases MAP2K1/2 was the most informative for predicting breast cancer cell sensitivity, and the predicted high-synergy combination of trametinib/omipalisib was validated experimentally in a PDX-derived TNBC cell line.

## 4. Methods

### Data Sources

The primary data sources we used can be split into three categories: the kinome profiling data set, the combination-treated cell line sensitivity set, and the cancer cell line transcriptomics set. The kinome profiling data set from the kinobeads assay was downloaded from the supplementary materials of Klaeger et al. 2017 [15]. For cancer cell line sensitivity to kinase inhibitor combinations, data was downloaded from (1) NCI-ALMANAC: cell sensitivity data was downloaded from the NCI wiki database (https://wiki.nci.nih.gov/display/NCIDTPdata/NCI-ALMANAC) and (2) Supplementary materials of previous lab combination screens published in Beville et al. 2019 [28] and Stuhlmiller et al. 2015 [29]. The CCLE gene expression set (“CCLE_expression.csv”) was downloaded from the DepMap portal (https://depmap.org/portal/download/all/) to create the set of cancer cell lines and their gene expression characteristics. In-house baseline gene expression data for the PDX-derived WHIM12 line was downloaded from the GEO repository for the Zawitowski et al. paper[26] (GSE87424).

### Data Preprocessing

The scripts implementing these descriptions are all available through github.

#### Klaeger et al. Kinobead Kinase Inhibition Profiles

As previously described [16], we read the values from the supplemental data table into R and produced a filtered list of kinase and kinase interactor relative intensity values. We imputed missing values with the default “no interaction” value of 1 and truncated likely outlier values to the 99.99 percentile (3.43).

#### Creating the Combination Inhibition State Data Set

To create a “combined” inhibition state of a given kinase inhibitor combination, we sought to superimpose the inhibition states of two individual states at specific doses. There were eight doses measured for each individual inhibitor, thus there were 64 possible combinations for each combination. We took the monotherapy kinome inhibition states from the Klaeger et al. set and computed a “combined” inhibition state for each kinase, based on three different combination schemes:

1. Simple Multiplicative: The simple conditional probability rule assumes two independent events (Eq. 1). Since the default “no interaction” inhibition value is 1, for kinases that are not targeted by both inhibitors simultaneously, the “combined” inhibition state value is simply either one in monotherapy.
2. Truncated Multiplicative: A minority of measured kinase inhibition states (∼1%) have values > 1 in the Klaeger et al. dataset, a possible artifact from the mass spectrometry measuring process. To avoid inflating those values, all >1 values were truncated at 1 and simple multiplication was performed as described above.
3. Addition: All kinase inhibition states were inverted into “Percent Inhibition” values, where 0 denotes no inhibition and 100 denotes complete inhibition. Then, when two inhibition states were combined, they were simply added together (truncated at a max value of 100).

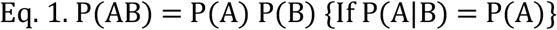

All three methods were tested in downstream modeling, resulting in minor variation. Truncated multiplied vectors were slightly more predictive (R^2^ score of ∼0.01 greater) so we used that scheme for all downstream modeling. In this way, we were able to compute hypothetical “combined” inhibition states for all possible combinations of ∼220 inhibitors, altogether comprising ∼2,000,000 combined inhibition states.

#### Dataset of Cancer Cell Line Sensitivity to Kinase Inhibitor Combinations

The cell sensitivity dataset from NCI-ALMANAC and previous lab publications were filtered to contain only kinase inhibitor small molecules, then summarized over replicates and converted to cell viability (1 = fully viable cell and 0 = full cell death). Relevant cancer types were annotated and individual cancer type datasets were subsetted for downstream cancer type-specific modeling.

#### Matching of Kinase Inhibitors between Inhibition State Dataset and Cell Line Sensitivity Dataset

The drug names from each dataset were read into R, and the package Webchem [30] was used to retrieve PubChem compound IDs (cid’s). The two sets of drug names were then matched based on these reference IDs, with a total of ∼100 matches between the two sets.

#### Baseline Gene Expression from CCLE

Data was preprocessed as described before [31] from the “CCLE_expression.csv” file. Cell line names were matched manually between CCLE and the NCI naming scheme. All cell lines represented in NCI-ALMANAC had a match in the CCLE database.

#### String

he STRING database [32] was processed as described previously [31] to annotate kinases and kinase interacting genes.Modeling Techniques. To assess our models we used a random 5-fold cross validation strategy. The features included in each fold were selected as specified by the feature selection scheme described in the results section. We implemented Elastic-net regression using the glmnet engine [33] for the feature selection scheme [17], We compared the performance of three model types using this strategy: random forest using the ranger engine [34] and gradient boosting using the XGBoost (eXtreme Gradient Boosting) engine [35]. Model performance was assessed by the R-squared value between predicted and actual outcome within the cross-validation scheme. For each model type and for the feature selection model, we tuned sets of 20 hyperparameters to find the best possible performer as follows: (a) Elastic-net: Penalty (0 - 0.1), Regularization (0.1-1) (b) Random Forest: Trees (100 - 2000) (c) XGBoost: Trees (100 - 1000), Tree Depth (4 - 30). After final model selection, we fit the model on the entire dataset and then made predictions on the experimental validation data.

All of the code written to support this paper is available through github (https://github.com/gomezlab/kinotype_combination_prediction) along with a brief walkthrough explaining where to find the code relevant to each part of the paper.

### Experimental Validation

6x6 dose combination screens were performed in the WHIM12 cell line as described in Beville et al. 2019 [28]. Briefly, cells were seeded in 384-well plates and dosed with drug after 24h. The screening library was tested for growth inhibition alone or in combination with Trametinib across 6 doses: 10 nmol/L, 100 nmol/L, 300 nmol/L, 1 μmol/L, 3 μmol/L, and 10 μmol/L. 0.1% DMSO was included as the control for growth inhibition on each plate. Plates were incubated at 37°C for 96 hours and lysed using CellTiter-Glo Reagent (Promega, catalog. no. G7570). Luminescence was measured using a PHERAstar FS instrument and growth inhibition was calculated relative to DMSO-treated wells.

## 5. Acknowledgements

We would like to thank UNC Research Computing for access to the computational resources necessary for this work. We would like to thank Michael P. East for his help with data compilation. This work was supported by grants through the National Institutes of Health (Grant #s CA274298, CA233811, CA238475, DK116204)

This is a preprint of an article submitted for consideration in Pacific Symposium on Biocomputing © 2024 [copyright World Scientific Publishing Company] [psb.stanford.edu]

## Notes

### Competing Interest Statement

The authors have declared no competing interest.

